# Induced systemic resistance by the root colonization of *Trichoderma atroviride* is independent from the chitin-mediated signaling pathway in Arabidopsis

**DOI:** 10.1101/2024.05.03.592335

**Authors:** Ayae Sakai, Hisako Yamagata, Keigo Naito, Takaya Tominaga, Shinsuke Ifuku, Hironori Kaminaka

**Affiliations:** Department of Agricultural Science, Graduate School of Sustainable Science, Tottori University, Tottori 680-8553, Japan; The United Graduate School of Agricultural Science, Tottori University, Tottori 680-8553, Japan; Graduate School of Engineering, Tottori University, Tottori 680-8552, Japan; Unused Bioresource Utilization Center, Tottori University, Tottori, 680-8550, Japan; Faculty of Agriculture, Tottori University, Tottori 680-8553, Japan

**Keywords:** Arabidopsis thaliana, *Alternaria brassicicola*, chitin, induced systemic resistance, *Trichoderma*

## Abstract

Beneficial root endophytic fungi induce systemic responses, growth promotion, and induced systemic resistance (ISR) in the colonized host plants. Soil application of chitin, a main component of fungal cell walls, also systemically induces disease resistance. Thus, chitin recognition and its downstream signaling pathway are supposed to mediate ISR triggered by beneficial fungi colonizing the root. This study compared systemic disease resistance and transcriptional responses induced by *Trichoderma*, a representative beneficial root endophytic fungus, and chitin in Arabidopsis. A significant plant growth promotion was observed under root colonization by the three tested beneficial fungi, *Trichoderma atroviride*, *Serendipita indica*, and *S. vermifera*. Still, only *T. atroviride* and *S. indica* triggered ISR against the necrotrophic fungal pathogen *Alternaria brassicicola*. Induced systemic resistance triggered by *T. atroviride* was compromised in the chitin-receptor mutant, while systemic resistance caused by soil application of chitin was not. Transcriptome analysis demonstrated that the chitin-regulated genes are mostly shared with those regulated by *T. atroviride*, but many of the latter were specific. However, the commonly enriched gene ontologies for those regulated genes indicated that *T. atroviride* inoculation and chitin application systemically control similar transcriptional responses, mainly associated with cell wall functions. Taken together, *Trichoderma* could trigger ISR primarily independently from the chitin-mediated signaling pathway; however, chitin and *Trichoderma* would systemically induce similar cellular functions in ISR aboveground.

## Introduction

Plants have evolved a complex immune system against microbial pathogen infection (Jones and Dangl, 2006). For the surveillance of microbes in host extracellular spaces, they recognize conserved microbial elicitors called pathogen-/microbe-associated molecular patterns (PAMPs/ MAMPs) through pattern recognition receptors (PRRs) localized on the cell surface (Dodds and Rathjen, 2010). Well-studied PAMP/MAMP-PRR combinations include the flagellin epitope flg22-FLS2 (FLAGELLIN SENSITIVE2), leucine-rich-repeat-type PRR, for bacteria, and chitin-CERK1 (Chitin Elicitor Receptor Kinase 1), lysin-motif (LysM)-type PRR, for fungi in the model plant Arabidopsis (Shu *et al*., 2023). Perception of PAMPs/MAMPs leads to pattern-triggered immunity (PTI), which activates a cellular defense response to restrict microbial invasion (Yuan *et al*., 2021). The microbial pathogens render the plant susceptible to disease by deploying virulence effectors into host cells to suppress PTI. However, plants recognize pathogen effectors via intracellular nucleotide-binding leucine-rich repeat receptors and induce a strong defense response accompanied by hypersensitive cell death called effecter-triggered immunity (ETI) (Jones and Dangl, 2006).

This plant immune system comprises similar cell-autonomous events to the innate immunity in animals, but, unlike animals, plants lack the adaptive immune system (Dodds and Rathjen, 2010). Therefore, plants have also developed original systemic immune systems to induce disease resistance against the next pathogen attack in distal parts from the infection site (Pieterse *et al*., 2014). Systemic acquired resistance (SAR) is a well-studied systemic immunity triggered by PTI and ETI upon pathogen infection. Induction of SAR depends on salicylic acid (SA) and is a long-lasting form of disease resistance against a broad spectrum of (hemi-)biotrophic pathogens (Durrant and Dong, 2004; Vlot *et al*., 2021). On the other hand, systemic immunity can also be triggered by beneficial or commensal microbes in the plant’s rhizosphere; it is termed induced systemic resistance (ISR) (Pieterse *et al*., 2014; Vlot *et al*., 2021). Unlike SAR, ISR depends on jasmonic acid and ethylene (ET), which function antagonistically to SA and act mainly against necrotrophic pathogens. Root endophytes include plant growth-promoting rhizobacteria (PGPR), such as *Pseudomonas* spp. and *Bacillus* spp., and plant growth-promoting fungi (PGPF), including *Trichoderma* spp. and *Serendipita* spp.; they are known as ISR-inducing rhizospheric microbes (Barazani *et al*., 2007; Ray *et al*., 2018; Vlot *et al*., 2021). As their names suggest, PGPR and PGPF can promote plant growth through root colonization. Additionally, root colonization by arbuscular mycorrhizal (AM) fungi, which establish a mutual symbiosis with approximately 70% of terrestrial plants by exchanging photosynthates for soil-derived mineral nutrients, also triggers ISR (Cameron *et al*., 2013).

Chitin is a β-1,4-linked linear polymer of *N*-acetylglucosamine and a well-known elicitor derived from fungal cell walls that induces disease resistance (Sharp, 2013). Additionally, soil application of chitin improves plant growth in various crops, which is thought to be independent of induced disease resistance. We have recently reported that supplementing soils with chitin systemically induces disease resistance against necrotrophic fungal pathogens in Arabidopsis, cabbage, strawberry, and rice (Parada *et al*., 2018; Takagi *et al*., 2022). Thus, chitin application to soils and beneficial root endophytic fungi induce similar systemic responses in plants, growth promotion and disease resistance. This similarity infers that ISR by beneficial fungi occurs through chitin recognition and its downstream signaling pathway. This study compared systemic disease resistance induced by a representative PGPF, *Trichoderma*, and chitin against a necrotrophic fungal pathogen, *Alternaria brassicicola*, in *Arabidopsis thaliana*. The evaluation of systemic disease resistance using a chitin-receptor CERK1 mutant and transcriptome analysis demonstrated that *Trichoderma* induces systemic disease resistance primarily independently from the chitin-mediated signaling pathway.

## Materials and Methods

### Plant and fungal materials

*Arabidopsis thaliana* (L) Heynh. accession Col-0 and *cerk1-2* (GABI_096F06) (Miya *et al*., 2007) were used as the wild-type and chitin-receptor mutant, respectively. *Trichoderma atroviride* ATCC 20476 and *Serendipita vermifera* MAFF305830 (Warcup, 1988) were maintained on potato dextrose agar (PDA: Difco, NJ, USA) medium at 25°C. *Serendipita indica* WP2 (Sherameti *et al*., 2005) was maintained on 1/6-strength Czapek Dox agar medium containing 0.8 g/L yeast extract and 15 g/L agar at 25°C.

### Plant growth conditions, fungal inoculation, and chitin application

Arabidopsis seeds were sown on a sterilized culture soil (Bestmix No. 3; Nippon Rockwool, Japan) and grown under controlled environmental conditions with 14-h light/8-h dark cycles at 23°C for seven weeks by watering the 1000-fold diluted fertilizer (HYPONeX [N–P–K = 6–10–5]; Hyponex Japan, Osaka, Japan) weekly.

For fungal inoculation, pieces of agar medium plugs of maintained fungal strains were placed on YEPG medium (Yeast extract 3 g/L, Polypepton 3 g/L, D-glucose 20 g/L) and cultured using a rotary shaker at 25°C under dark condition for two weeks. After removing the liquid medium, the harvested mycelium was homogenized with a blender (Nissei, Osaka, Japan) at 10,000 rpm for 10 sec. Distilled water was added to prepare the fungal suspension at the indicated concentrations. Five mL of each fungal suspension was irrigated into soils two weeks after sowing, and Arabidopsis seedlings were grown for an additional five weeks. The water dispersion of chitin nanofiber (CNF) (MARINE NANO-FIBER CN-01), which is produced directly from chitin powder by physically grinding the microfibrils (nanofibrillation) and can be used like a water solution (Ifuku and Saimoto, 2012), was purchased from Marine Nano-fiber (Tottori, Japan) and used for chitin treatment. Upon preparation, the culture soil was mixed with an equal volume of 0.1% (w/v) CNF water dispersion before sowing, based on our previous study (Kaminaka *et al*., 2020). Distilled water was used for control experiments.

### Fluorescent staining of fungal hyphae in Arabidopsis roots

The harvested Arabidopsis roots were fixed in 70% ethanol overnight. After removing ethanol, 5% KOH was added, and samples were heated at 90°C for 30 min. Root samples were transferred into 1% HCl for neutralization for 5 min, washed with PBS buffer, and incubated with 5 μg/mL of WGA-Alexa Fluor 488 (Thermo Fisher Scientific, MA, USA) under dark conditions for 10 min. Roots were washed with PBS again, and 20% TOMEI (Tokyo Chemical Industry, Tokyo, Japan) was added to clear tissues. The stained roots were observed under a fluorescent microscope (DM2500; Leica, Wetzlar, Germany) with an excitation filter L5, and photo images were obtained with the equipped digital camera (DFC310; Leica).

### Disease resistance assay

Spores of the fungal pathogen *A. brassicicola* strain O-264, a causal agent of black leaf spot of Brassica plants, were prepared and inoculated on Arabidopsis leaves according to a previous study (Parada *et al*., 2018). Ten μL of conidial suspension (5.0×10^5^ spores/mL) was inoculated on each leaf. The diameter of emerged lesions on leaves was measured four days after the inoculation using ImageJ Ver.1.53a.

### RNA-sequencing and data analysis

About 100 mg of randomly selected leaves from at least three individual seven-week-old Arabidopsis seedlings inoculated with *T. atroviride* or treated with chitin were used to prepare the sequencing library preparation. Total RNA preparation was conducted according to Tominaga *et al*. (2021). Preparation of sequencing libraries and sequencing with strand-specific and paired-end reads (150 bp) by DNBSEQ-T7RS was performed by Genome-Lead Co. (Kagawa, Japan).

Low-quality reads (< QV30) and adapter sequences of the obtained raw reads were removed by fastp (Chen *et al*., 2018) and mapped onto the sequence of the Arabidopsis reference genome TAIR10.43 (https://www.arabidopsis.org/) by an RNA-sequencing aligner STAR Ver.2.6.1d (Dobin *et al*., 2013) (Supplementary Table S1). The data were processed with featureCounts Ver.2.0.1 (Liao *et al*., 2014) to obtain the gene expression count data. Each count data in different library sizes was normalized by the trimmed mean of the *M*-values normalization method, and differentially expressed genes (DEGs) were identified by comparing control and *T. atroviride*-inoculated or chitin-treated plants using edgeR Ver.4.2.1 (Robinson *et al*., 2010). The list of DEGs was extracted by a false discovery rate (FDR) cutoff < 0.05. Venn diagrams were generated using the Venn diagram website (https://bioinformatics.psb.ugent.be/webtools/Venn). The gene ontology (GO) enrichment analysis was conducted using Shiny GO 0.77 (Ge *et al*., 2020), and the dot plots were drawn using “ggplot2” and “ggpubr” packages in R (Ver.4.3.1).

### RNA-sequencing data accession number

The raw read data obtained by RNA-sequencing were deposited in the DNA Data Bank of Japan under the BioProject accession number PRJDB17932.

### Statistical analysis

Lesion diameters caused by *A. brassicicola* inoculation were compared to control plants, and the statistical significance of the results was analyzed using Student’s *t*-test and Microsoft Excel (Ver.2312). All pathogen inoculation tests were conducted at least three times with more than three different plants for each treatment and genotype.

## Results

### Growth promotion and ISR by the beneficial fungi colonization of Arabidopsis

Root endophytes promote plant growth and systemically induce disease resistance; they are termed PGPR and PGPF (Pieterse *et al*., 2014; Vlot *et al*., 2021). In Arabidopsis, *Trichoderma* and *Serendipita* species are known as PGPF (Lahrmann and Zuccaro, 2012; González-Pérez *et al*., 2018). First, to compare the effects of beneficial fungi inoculated on Arabidopsis seedlings, *T. atroviride*, *S. indica,* and *S. vermifera* were inoculated by irrigating the soil with fungal suspensions. The growth of Arabidopsis seedlings inoculated with these three fungi was significantly promoted compared to non-inoculated seedlings at five weeks post-inoculation (Fig. 1A). All fungal colonization was confirmed by hyphal staining only in the roots of fungus-inoculated seedlings (Fig. 1B). These results indicate that root colonization by these beneficial fungi promotes plant growth.

**Fig. 1.**
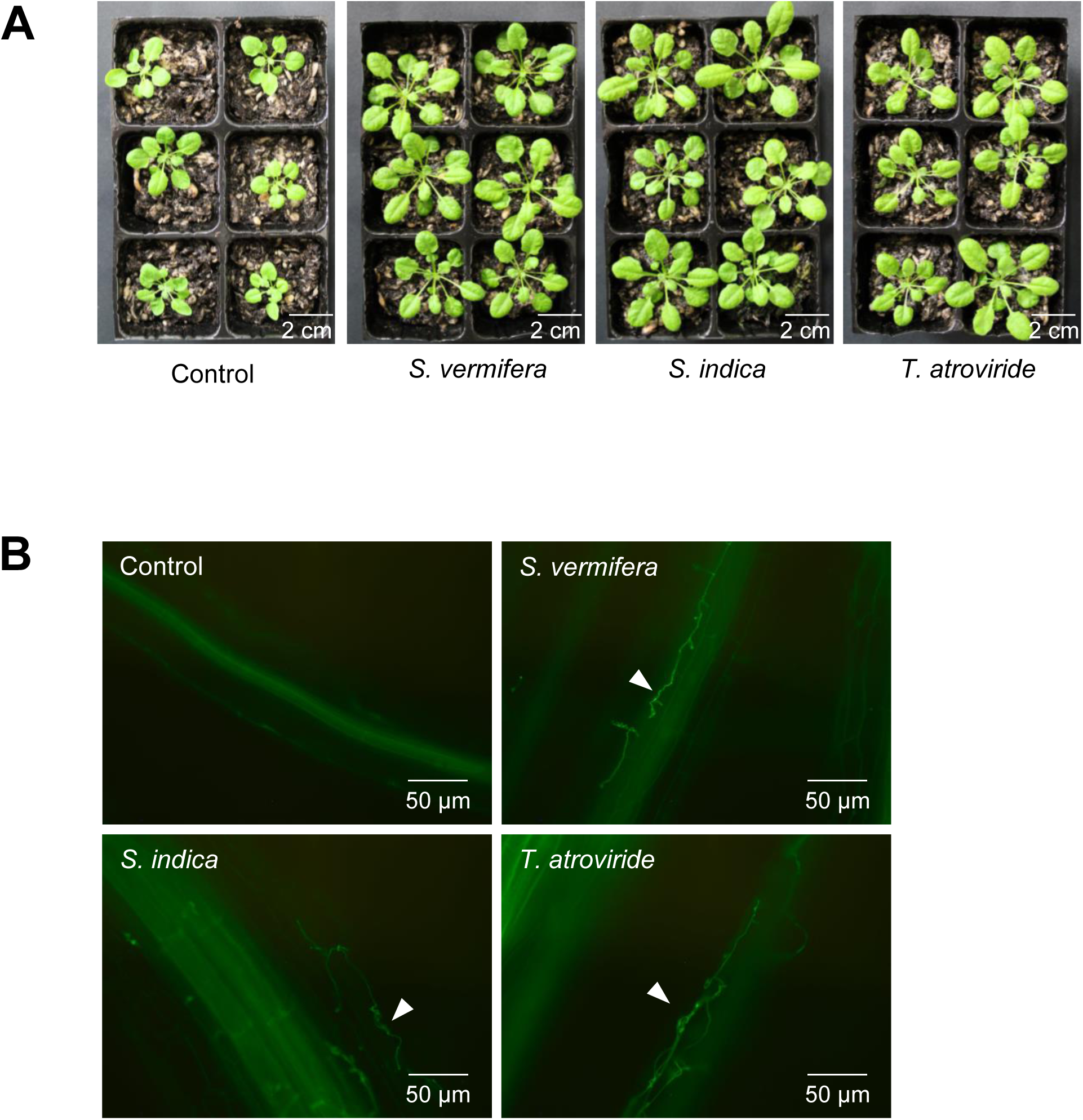
Growth promotion of Arabidopsis seedlings colonized by beneficial fungi. (A) Photos of Arabidopsis seedlings grown for seven weeks on soil irrigated with hyphal homogenates (50 mg fresh weight [FW]/mL) of *Serendipita vermifera*, *S. indica*, and *Trichoderma atroviride*. (B) Fluorescent images of inoculated fungal hyphae stained with WGA-Alexa Fluor 488 in Arabidopsis roots. The white arrowheads indicate fungal hyphae.

Since beneficial fungi cause ISR mainly against necrotrophic pathogens (Pieterse *et al*., 2014; Vlot *et al*., 2021), we examined disease resistance against the necrotrophic fungal pathogen *A. brassicicola*, a causal agent of black leaf spot of Brassica plants, in the leaves of beneficial fungi-inoculated seedlings. Root colonization by *T. atroviride* and *S. indica* significantly reduced the lesion size. In contrast, no significant differences in lesion formation were observed in *S. vermifera*-colonized seedlings compared to the control experiment (Fig. 2A). Since *T. atroviride* displayed significantly more ISR compared to *S. indica*, it was chosen for the following experiment to optimize the fungal inoculum concentration. Only the concentration used for the previous experiment (50 mg FW/mL), not lower ones, led to significantly reduced lesions caused by *A. brassicicola* infection (Fig. 2B). Thus, that concentration of *T. atroviride* inoculum was used for further experiments.

**Fig. 2.**
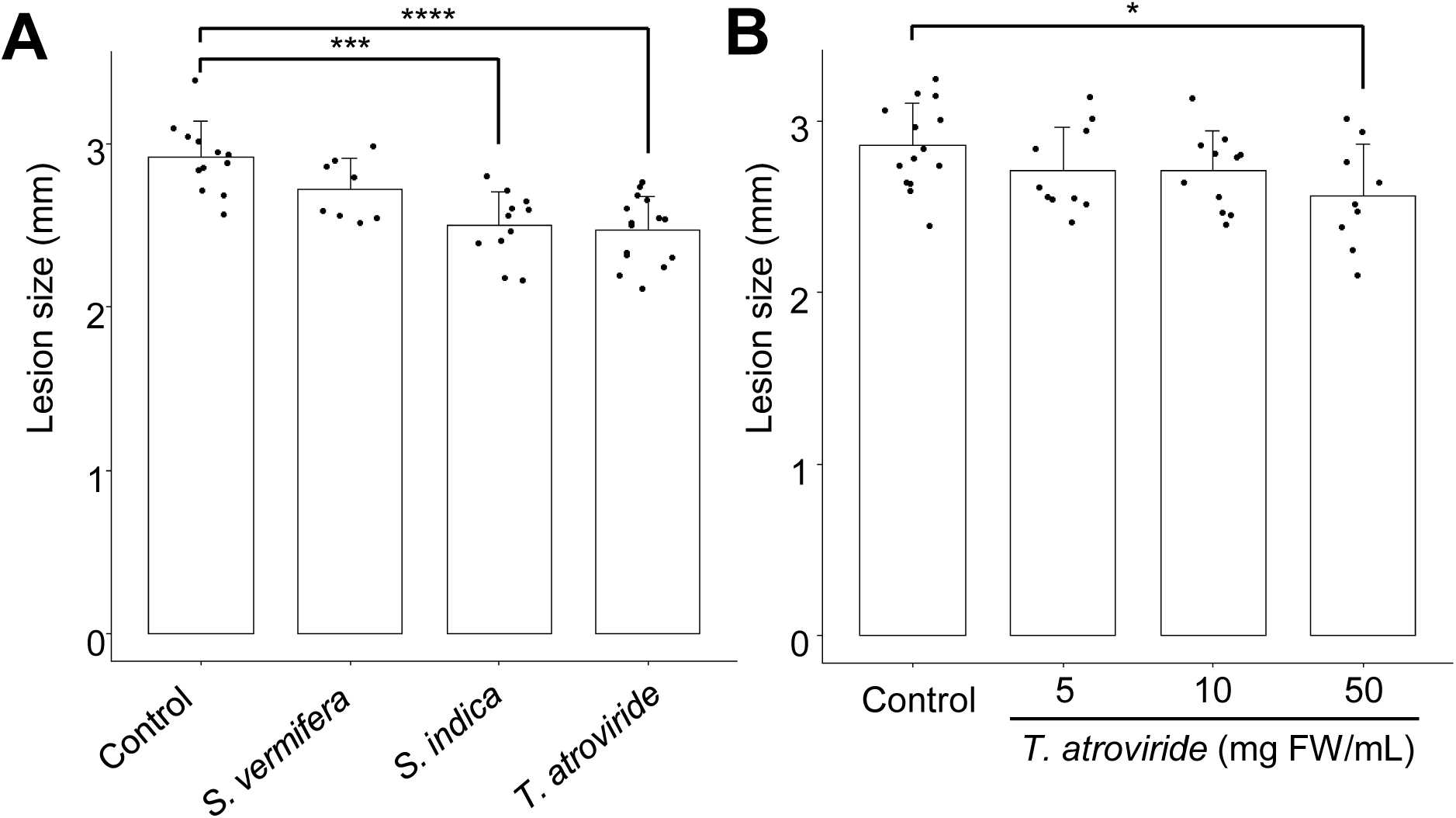
Induced systemic resistance against necrotrophic pathogen after inoculation with beneficial fungi. (A) Lesion size on Arabidopsis seedling leaves (grown as in Fig. 1) inoculated with 10 μL of *Alternaria brassicicola* suspension (5.0×10^5^ spores/mL). The lesion was measured four days after inoculation. (B) The effects of inoculum concentration (mg fresh weight [FW]/mL) for *Trichoderma atroviride* inoculation conducted as in (A). The bars and error bars indicate mean and standard errors, and asterisks indicate statistically significant differences (Student’s *t*-test: \**P* < 0.05, \*\*\**P* < 0.001, \*\*\*\**P* < 0.0001; n ≥ 8).

### Chitin receptor CERK1 function in ISR by T. atroviride and chitin

Chitin is a PAMP/MAMP used by plants to sense the presence of fungi in intracellular spaces (Shu *et al*., 2023). Like beneficial fungus colonization, applying chitin into soils promotes plant growth and induces systemic resistance (Parada *et al*., 2018; Kaminaka *et al*., 2020; Takagi *et al*., 2022). Therefore, the involvement of chitin in the ISR caused by *T. atroviride* root colonization was examined using the chitin receptor LysM-type PRR CERK1 deficient mutant *cerk1-2* (Miya *et al*., 2007). Chitin nanofibers were used as chitin in this study because they can induce a more robust plant response than other chitins (Egusa *et al*., 2015; Kaminaka *et al*., 2020). The ISR against *A. brassicicola* observed in wild-type plants inoculated with *T. atroviride* was compromised in *cerk1-2* (Fig. 3A). In contrast, the systemic disease resistance against *A. brassicicola* was significantly induced even in *cerk1-2* by chitin application into soils (Fig. 3B). These distinct results reveal that the chitin-triggered function does not contributes to ISR caused by *T. atroviride* root colonization of Arabidopsis.

**Fig. 3.**
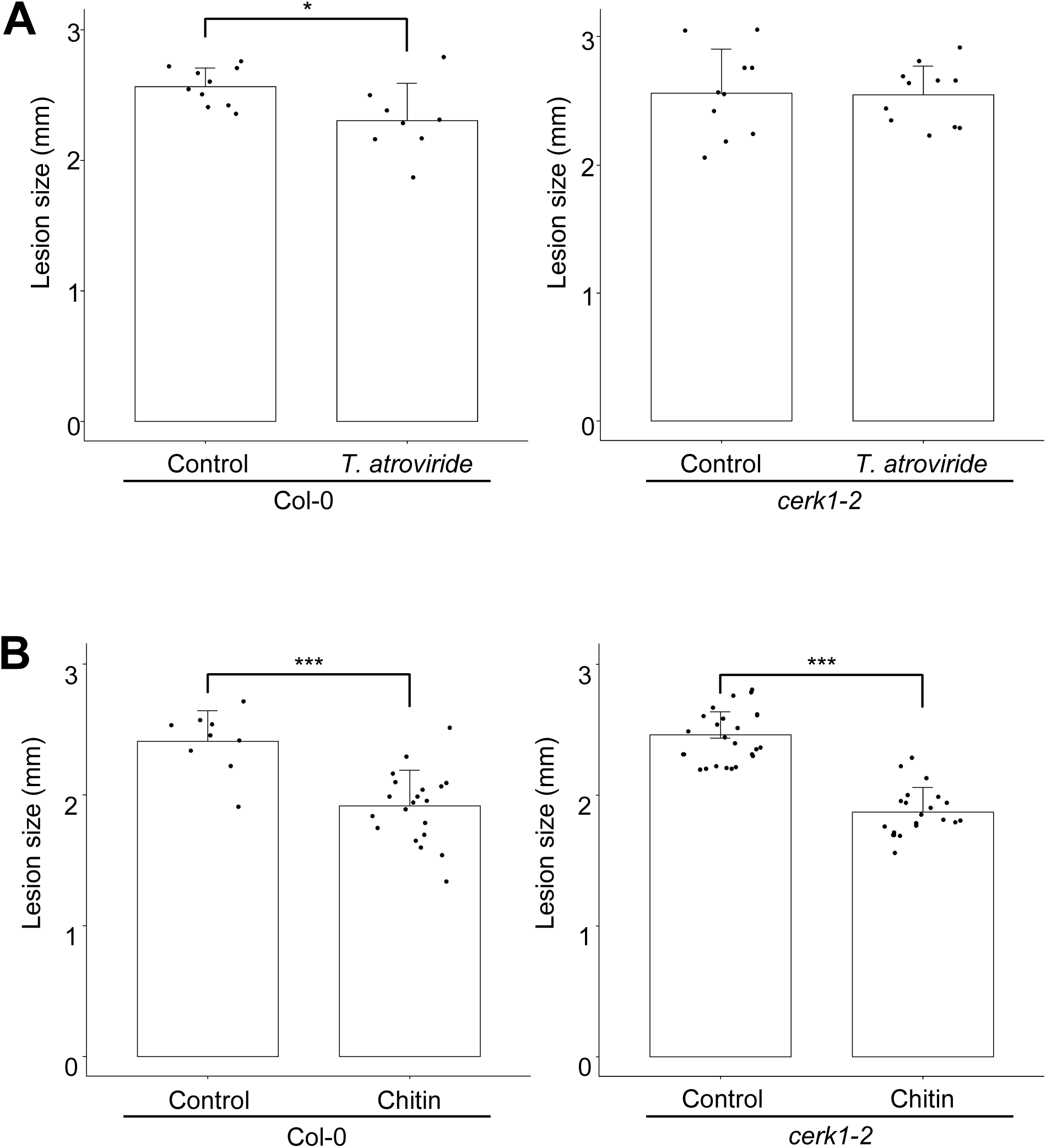
Effects of chitin receptor deficiency on induced systemic resistance against a necrotrophic pathogen. Disease resistance against *Alternaria brassicicola* on leaves of wild-type (Col-0) and *cerk1-2* seedlings (A) inoculated with *Trichoderma atroviride* and (B) treated with chitin, as conducted in Fig. 2. The bars and error bars indicate mean and standard errors, and asterisks indicate statistically significant differences (Student’s *t*-test: \**P* < 0.05, \*\*\**P* < 0.001; n ≥ 8).

### Comparative transcriptome analysis of T. atroviride-inoculated or chitin-treated Arabidopsis seedlings

To elucidate the molecular mechanism underlying ISR caused by the root colonization of *T. atroviride*, Arabidopsis roots were inoculated with *T. atroviride* or treated with chitin, and seedling leaves were used for RNA-sequencing to elucidate the molecular mechanism underlying ISR caused by *T. atroviride*. Compared with control seedlings, 1,724 DEGs were identified in *T. atroviride*-inoculated seedlings, including 617 upregulated and 1,107 downregulated DEGs (Fig. 4A, Supplementary Table S2). We identified 95 DEGs in chitin-treated seedlings, including 24 upregulated and 71 downregulated DEGs (Fig. 4B, Supplementary Table S3). Notably, more than 95% of DEGs in chitin-treated seedlings were shared with those in *T. atroviride*-inoculated seedlings (Fig. 4A, B).

**Fig. 4.**
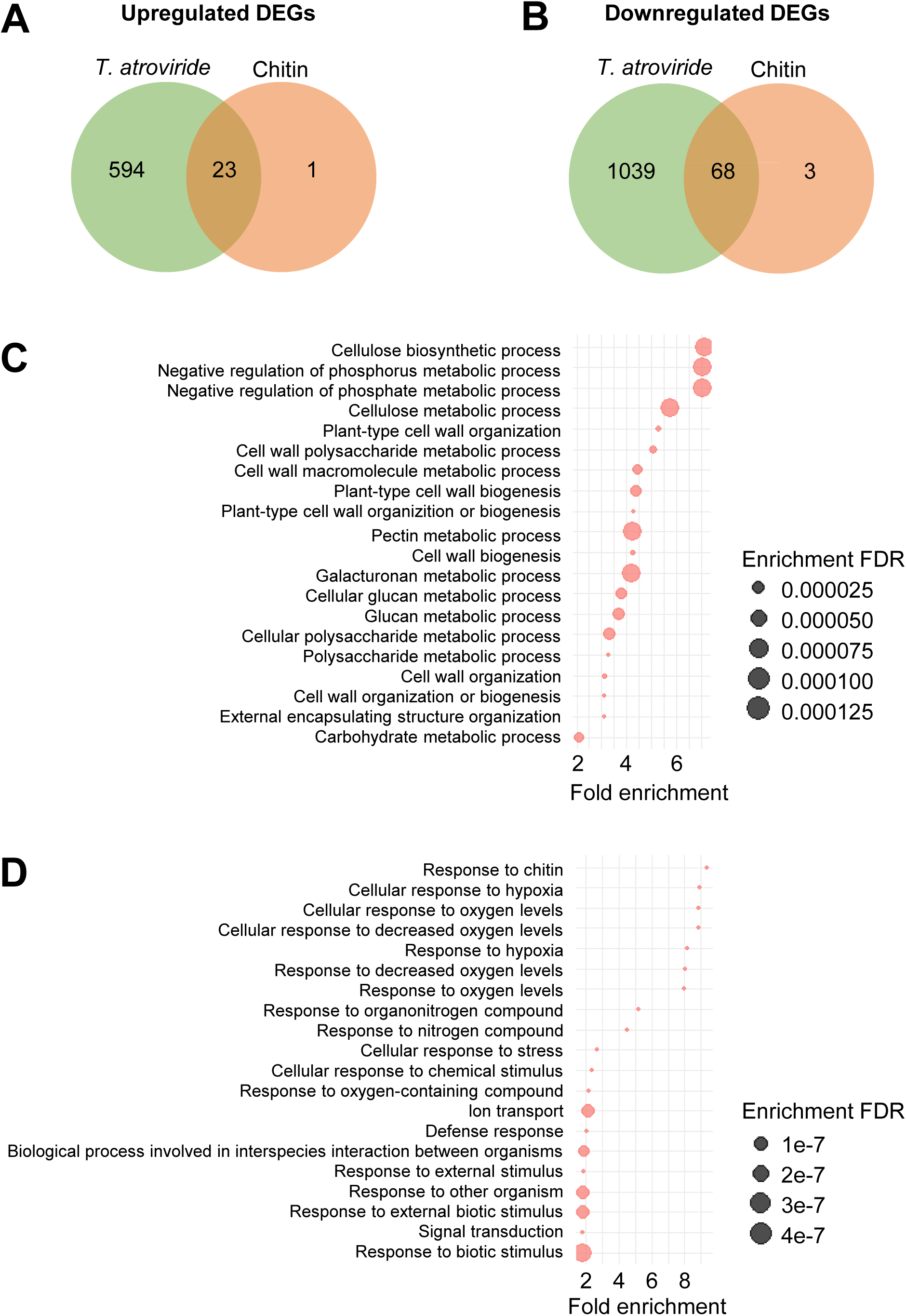
Transcriptome analysis of the leaves of Arabidopsis seedlings inoculated with *Trichoderma atroviride* and treated with chitin. Venn diagram of upregulated (A) and downregulated (B) differentially expressed genes (DEGs) identified by a false discovery rate (FDR) cutoff < 0.05. The top 20 enriched biological process gene ontology (GO) terms with the lowest FDR values for upregulated (C) and downregulated (D) DEGs upon *T. atroviride* inoculation. The circle size indicates the FDR value. The complete list of enriched GO terms is presented in Supplementary Table S4.

Next, GO enrichment analysis was conducted for DEGs identified in the leaves of *T. atroviride*-inoculated seedlings. Regarding upregulated DEGs, GO terms associated with cell wall function (e.g., “Cellulose biosynthesis/metabolic process,” “Plant-type cell wall organization or biogenesis,” and “Cell wall polysaccharide/macromolecule metabolic process”) were dominantly overrepresented in the biological process category (Fig. 4C, Supplementary Table S4) and were also found as enriched GO terms for upregulated DEGs in chitin-treated seedling leaves (Supplementary Fig. S1A, Table S5). The enrichment for categories related to the negative regulation of phosphorus metabolic process, kinase inhibitor activity, and endocytic pathway (e.g., endosome, Golgi, or vesicle) was also indicated (Fig. 4C, Supplemental Table S4). Regarding downregulated DEGs, the GO term “Response to chitin” was highly enriched, and overrepresented GO terms associated with cellular responses to oxygen levels, transcription factors, and NAD/NAD(P)+ nucleoside activity were also found (Fig. 4D, Supplementary Table S4). These GO terms were also overrepresented for downregulated DEGs in chitin-treated seedlings (Supplementary Fig. S1B, Table S5).

## Discussion

*Trichoderma* is widely used as a biocontrol agent mainly against soil-borne diseases in various crops through its mycoparasitism and secretomes, including volatile organic compounds (VOCs), cell wall-degrading enzymes (CDWEs), reactive oxygen species, and antimicrobial secondary metabolites (Alfiky and Weisskopf, 2021; Yao *et al*., 2023). Additionally, *Trichoderma* induces systemic responses in host plants, including growth promotion and disease resistance known as ISR. Combining these functions would cause the biocontrol effects of *Trichoderma*, but available knowledge on each function needs to be improved. To obtain novel insights into plant–*Trichoderma* interactions, we investigated the molecular mechanism underlying *Trichoderma*-induced systemic resistance by focusing on the involvement of chitin—recognition and signaling—in Arabidopsis. Analyzing systemic disease resistance in chitin receptor-deficient mutants and transcriptomes has revealed that *Trichoderma* induces systemic resistance against necrotrophic pathogens primarily independently from the chitin-mediated signaling pathway.

The root colonization of all examined endophytic fungi, *T. atroviride*, *S. indica*, and *S. vermifera*, promoted plant growth, consistent with findings in various plants (Lahrmann and Zuccaro, 2012; Ray *et al*., 2018; Alfiky and Weisskopf, 2021). In contrast, only *T. atrovirid*e and *S. indica* significantly increased the disease resistance against a necrotrophic pathogen, *A. brassicicola*, in the leaves of root-colonized Arabidopsis seedlings. The ISR by root colonization of *T. atroviride* and *S. indicia* in Arabidopsis is reported (Salas-Marina *et al*., 2011; Lahrmann and Zuccaro, 2012), but not for *S. vermifera.* Sarkar *et al*. (2019) have revealed the mycoparasitism of *S. vermifera* against *Bipolaris sorokiniana*, a causal agent of spot blotch and common root rot diseases, by mainly reducing the pathogen’s root infection in barley. They also suggested a disease resistance systemically induced by *S. vermifera* in roots but with no statistical significance. Thus, *S. vermifera* may be able to cause weak ISR, which would not be enough to be confirmed using our Arabidopsis pathosystem.

In Arabidopsis, ISR has been well-studied using PGPR such as *Pseudomonas* and *Bacillus* species (Pieterse *et al*., 2014; Vlot *et al*., 2021). However, knowledge of ISR by beneficial fungi is limited due to AM fungi being non-host. Our study demonstrated that the function of the chitin-receptor CERK1 is required for ISR by *Trichoderma* in Arabidopsis. The ectomycorrhizal fungus *Laccaria bicolor* triggered ISR against the cabbage looper *Trichoplusia ni* and induced systemic susceptibility against the hemi-biotrophic bacterial pathogen *Pseudomonas syringae* pv. *tomato* DC3000 in non-host Arabidopsis plants in a CERK1-dependent way (Vishwanathan *et al*., 2020). Since the treatments of heat-killed *L. bicolor* and chitin also systemically induced disease resistance, the authors proposed that ISR without symbiotic association would be triggered by the root perception of PAMPs/MAMPs, which is supported by our previous study (Takagi *et al*., 2022). In rice, supplementing soils with chitin systemically induced disease resistance against the necrotrophic pathogen *Bipolaris oryzae* through the function of LysM-type PRRs, OsCERK1, and OsCEBiP. In contrast, chitin-induced systemic resistance in Arabidopsis was not mediated by CERK1 in this study. The mechanism for chitin perception by LysM-type PRRs in Arabidopsis is similar to that in rice, but CERK1 function is different in terms of its binding ability to chitin: unlike rice CERK1 (OsCERK1), Arabidopsis CERK1 can bind to chitin oligosaccharides (Yang *et al*., 2022). Thus, the difference in the chitin perception mechanism may be explained by the different requirements of CERK1 function for chitin-induced systemic resistance between Arabidopsis and rice. To address this point, Arabidopsis LysM-type PRR(s) involved in chitin-induced systemic resistance and *Trichoderma*-induced ISR should be characterized using loss-of-function mutants, which will be conducted in a subsequent study.

The transcriptome analysis revealed that most DEGs identified in chitin-treated seedlings were shared with those in *Trichoderma*-inoculated seedlings, indicating the minor or no contribution of chitin-triggered functions in *Trichoderma*-induced ISR. Additionally, 94% of DEGs identified in *Trichoderma*-inoculated seedlings were specific; therefore, *Trichoderma*-specific transcriptional responses would contribute to ISR. The GO terms involved in cell wall functions were mainly overrepresented by the GO enrichment analysis of upregulated DEGs in *Trichoderma*-inoculated seedlings. Recently, many reports revealed the involvement and importance of cell wall functions (e.g., cell wall biogenesis, composition, and integrity) in inducing disease resistance (Bacete *et al*., 2018; Molina *et al*., 2021; Baez *et al*., 2022). Thus, the modulation of cell wall conditions by transcriptional changes would be a major cellular event inducing disease resistance aboveground in *Trichoderma*-triggered ISR. In our previous study, the GO terms associated with cell wall functions were also enriched in the leaves of rice seedlings grown on chitin-supplemented soils (Takagi *et al*., 2022). However, unlike in Arabidopsis, the genes involved in cell wall functions were downregulated in rice. These opposite aboveground transcriptional responses may also describe the different requirements of CERK1 function for chitin-induced systemic resistance between Arabidopsis and rice.

The enriched GO terms for DEGs were similar between *Trichoderma* inoculation and chitin treatment. Thus, this inoculation/treatment would systemically induce disease resistance by modulating similar cellular functions mainly associated with aboveground cell wall function, even if the requirement of LysM-type PRRs is quite different (Fig. 5). In this study, we identified a specific signaling pathway mediated by CERK1 in ISR by *Trichoderma*, which is primarily independent of the chitin-mediated signaling pathway. Plant hormones and secretomes, including effectors, VOCs, and CDWEs, also participate in ISR by *Trichoderma* (Alfiky and Weisskopf, 2021). Therefore, we plan to conduct further studies focusing on the involvement of these molecules to elucidate the mechanism underlying the *Trichoderma*-specific signaling pathway to induce disease resistance in Arabidopsis systemically.

**Fig. 5.**
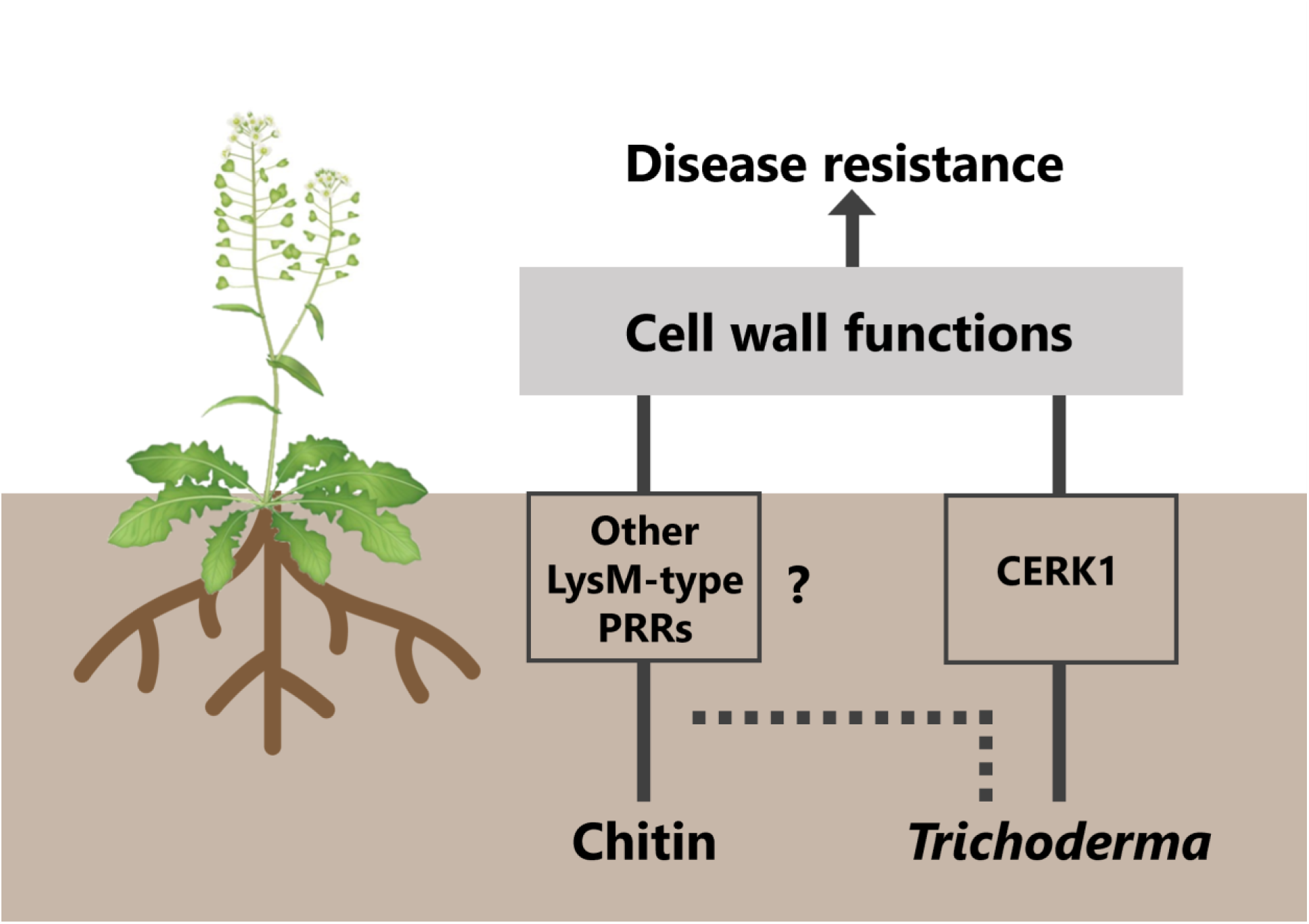
Proposed model for chitin- and *Trichoderma*-induced systemic resistance (ISR) in Arabidopsis. Chitin supplementation into soils and root colonization by *Trichoderma atroviride* systemically upregulate cell wall-related genes in leaves and induce disease resistance against the necrotrophic fungal pathogen *Alternaria brassicicola*. The function of CERK1 (Chitin Elicitor Receptor Kinase 1), lysin-motif (LysM)-type pattern recognition receptor required for chitin perception (Miya *et al*., 2007), is required for ISR by *Trichoderma*, whereas CERK1 is not involved in ISR by chitin. Hence, chitin and *Trichoderma* would systemically modulate similar cellular functions for aboveground ISR. However, *Trichoderma* induces systemic responses primarily independently from the chitin-mediated signaling pathway.

## Acknowledgments

We thank Dr. Hirofumi Nakagami (Max-Planck Research Institute), Dr. Hiroshi Otani (Tottori University), and Dr. Patrick Schäfer (Justus Liebig University) for providing *cerk1-2* seeds, *Alternaria brassicicola*, and *Serendipita indica*, respectively. This study was supported by the Japan Society for the Promotion of Science (JSPS) KAKENHI grant number 22K19182.

---

Supplementary Fig. S1. Enriched biological process GO terms for upregulated (A) and downregulated (B) DEGs identified by the transcriptome analysis of leaves of Arabidopsis seedlings treated with chitin. The circle size indicates the FDR value. The complete list of enriched GO terms is presented in Supplementary Table S5.

Supplementary Table S1. Summary of RNA-sequencing

Supplementary Table S2. List of DEGs identified in leaves of *T. atroviride*-inoculated Arabidopsis seedlings

Supplementary Table S3. List of DEGs identified in leaves of chitin-treated Arabidopsis seedlings

Supplementary Table S4. Enriched GO terms for DEGs identified in leaves of *T. atroviride*-inoculated Arabidopsis seedlings

Supplementary Table S5. Enriched GO terms for DEGs identified in leaves of chitin-treated Arabidopsis seedlings

## References

Alfiky, A., and Weisskopf, L. (2021) Deciphering *Trichoderma*–plant–pathogen interactions for better development of biocontrol applications. J Fungi 7: 1–18.

Bacete, L., Mélida, H., Miedes, E., and Molina, A. (2018) Plant cell wall-mediated immunity: cell wall changes trigger disease resistance responses. Plant J 93: 614–636.

Baez, L.A., Tichá, T., and Hamann, T. (2022) Cell wall integrity regulation across plant species. Plant Mol Biol 109: 483–504.

Barazani, O., von Dahl, C.C., and Baldwin, I.T. (2007) *Sebacina vermifera* promotes the growth and fitness of *Nicotiana attenuata* by inhibiting ethylene signaling. Plant Physiol 144: 1223–1232.

Cameron, D.D., Neal, A.L., van Wees, S.C.M., and Ton, J. (2013) Mycorrhiza-induced resistance: more than the sum of its parts? Trends Plant Sci 18: 539–545.

Chen, S., Zhou, Y., Chen, Y., and Gu, J. (2018) fastp: an ultra-fast all-in-one FASTQ preprocessor. Bioinformatics 34: i884–i890.

Dobin, A., Davis, C.A., Schlesinger, F., Drenkow, J., Zaleski, C., Jha, S., et al. (2013) STAR: Ultrafast universal RNA-seq aligner. Bioinform 29: 15–21.

Dodds, P.N., and Rathjen, J.P. (2010) Plant immunity: towards an integrated view of plant–pathogen interactions. Nat Rev Genet 11: 539–548.

Durrant, W.E., and Dong, X. (2004) Systemic acquired resistance. Annu Rev Phytopathol 42: 185– 209.

Egusa, M., Matsui, H., Urakami, T., Okuda, S., Ifuku, S., Nakagami, H., and Kaminaka, H. (2015) Chitin nanofiber elucidates the elicitor activity of polymeric chitin in plants. Front Plant Sci 6: 1098.

Ge, S.X., Jung, D., and Yao, R. (2020) ShinyGO: a graphical gene-set enrichment tool for animals and plants. Bioinformatics 36: 2628–2629.

González-Pérez, E., Ortega-Amaro, M.A., Salazar-Badillo, F.B., Bautista, E., Douterlungne, D., and Jiménez-Bremont, J.F. (2018) The Arabidopsis-Trichoderma interaction reveals that the fungal growth medium is an important factor in plant growth induction. Sci Rep 8: 16427.

Humphrey, T. V, Bonetta, D.T., and Goring, D.R. (2007) Sentinels at the wall: cell wall receptors and sensors. New Phytol 176: 7–21.

Ifuku, S., and Saimoto, H. (2012) Chitin nanofibers: preparations, modifications, and applications. Nanoscale 4: 3308–3318.

Jones, J.D.G., and Dangl, J.L. (2006) The plant immune system. Nature 444: 323–329.

Kaminaka, H., Miura, C., Isowa, Y., Tominaga, T., Gonnami, M., Egusa, M., and Ifuku, S. (2020) Nanofibrillation is an effective method to produce chitin derivatives for induction of plant responses in soybean. Plants 9: 1–11.

Lahrmann, U., and Zuccaro, A. (2012) *Opprimo ergo sum*—evasion and suppression in the root endophytic fungus *Piriformospora indica*. Mol Plant Microb Interact 25: 727–737.

Liao, Y., Smyth, G.K., and Shi, W. (2014) featureCounts: an efficient general purpose program for assigning sequence reads to genomic features. Bioinformatics 30: 923–930.

Miya, A., Albert, P., Shinya, T., Desaki, Y., Ichimura, K., Shirasu, K., et al. (2007) CERK1, a LysM receptor kinase, is essential for chitin elicitor signaling in *Arabidopsis*. Proc Natl Acad Sci USA 104: 19613–19618.

Molina, A., Miedes, E., Bacete, L., Rodríguez, T., Mélida, H., Denancé, N., et al. (2021) *Arabidopsis* cell wall composition determines disease resistance specificity and fitness. Proc Natl Acad Sci USA 118: e2010243118.

Parada, R.Y., Egusa, M., Aklog, Y.F., Miura, C., Ifuku, S., and Kaminaka, H. (2018) Optimization of nanofibrillation degree of chitin for induction of plant disease resistance: elicitor activity and systemic resistance induced by chitin nanofiber in cabbage and strawberry. Int J Biol Macromol 118: 2185–2192.

Pieterse, C.M.J., Zamioudis, C., Berendsen, R.L., Weller, D.M., Van Wees, S.C.M., and Bakker, P.A.H.M. (2014) Induced systemic resistance by beneficial microbes. Annu Rev Phytopathol 52: 347–375.

Ray, P., Chi, M.-H., Guo, Y., Chen, C., Adam, C., Kuo, A., et al. (2018) Genome sequence of the plant growth promoting fungus *Serendipita vermifera* subsp. *bescii*: the first native strain from North America. Phytobiomes J 2: 62–63.

Robinson, M.D., McCarthy, D.J., and Smyth, G.K. (2010) edgeR: a bioconductor package for differential expression analysis of digital gene expression data. Bioinformatics 26: 139–140.

Salas-Marina, M.A., Silva-Flores, M.A., Uresti-Rivera, E.E., Castro-Longoria, E., Herrera-Estrella, A., and Casas-Flores, S. (2011) Colonization of *Arabidopsis* roots by *Trichoderma atroviride* promotes growth and enhances systemic disease resistance through jasmonic acid/ethylene and salicylic acid pathways. Eur J Plant Pathol 131: 15–26.

Sarkar, D., Rovenich, H., Jeena, G., Nizam, S., Tissier, A., Balcke, G.U., et al. (2019) The inconspicuous gatekeeper: endophytic *Serendipita vermifera* acts as extended plant protection barrier in the rhizosphere. New Phytol 224: 886–901.

Sharp, R.G. (2013) A review of the applications of chitin and its derivatives in agriculture to modify plant-microbial interactions and improve crop yields. Agronomy 3: 757–793.

Sherameti, I., Shahollari, B., Venus, Y., Altschmied, L., Varma, A., and Oelmüller, R. (2005) The endophytic fungus *Piriformospora indica* stimulates the expression of nitrate reductase and the starch-degrading enzyme glucan-water dikinase in tobacco and *Arabidopsis* roots through a homeodomain transcription factor that binds to a conserved motif in their promoters. J Biol Chem 280: 26241–26247.

Shu, L.-J., Kahlon, P.S., and Ranf, S. (2023) The power of patterns: new insights into pattern-triggered immunity. New Phytol 240: 960–967.

Takagi, M., Hotamori, K., Naito, K., Matsukawa, S., Egusa, M., Nishizawa, Y., et al. (2022) Chitin-induced systemic disease resistance in rice requires both OsCERK1 and OsCEBiP and is mediated via perturbation of cell-wall biogenesis in leaves. Front Plant Sci 13: 1064628.

Tominaga, T., Miura, C., Sumigawa, Y., Hirose, Y., Yamaguchi, K., Shigenobu, S., et al. (2021) Conservation and diversity in gibberellin-mediated transcriptional responses among host plants forming distinct arbuscular mycorrhizal morphotypes. Front Plant Sci 12: 795695.

Vishwanathan, K., Zienkiewicz, K., Liu, Y., Janz, D., Feussner, I., Polle, A., and Haney, C.H. (2020) Ectomycorrhizal fungi induce systemic resistance against insects on a nonmycorrhizal plant in a CERK1-dependent manner. New Phytol 228: 728–740.

Vlot, A.C., Sales, J.H., Lenk, M., Bauer, K., Brambilla, A., Sommer, A., et al. (2021) Systemic propagation of immunity in plants. New Phytol 229: 1234–1250.

Wang, Y., Liu, Z., Hao, X., Wang, Z., Wang, Z., Liu, S., et al. (2023) Biodiversity of the beneficial soil-borne fungi steered by *Trichoderma*-amended biofertilizers stimulates plant production. NPJ Biofilms Microbiomes 9: 46.

Warcup, J.H. (1988) Mycorrhizal associations of isolates of *Sebacina vermifera*. New Phytol 110: 227–231.

Yang, C., Wang, E., and Liu, J. (2022) CERK1, more than a co-receptor in plant–microbe interactions. New Phytol 234: 1606–1613.

Yao, X., Guo, H., Zhang, K., Zhao, M., Ruan, J., and Chen, J. (2023) Trichoderma and its role in biological control of plant fungal and nematode disease. Front Microbiol 14: 1160551.

Yuan, M., Ngou, B.P.M., Ding, P., and Xin, X.-F. (2021) PTI-ETI crosstalk: an integrative view of plant immunity. Curr Opin Plant Biol 62: 102030.

Zin, N.A., and Badaluddin, N.A. (2020) Biological functions of *Trichoderma* spp. for agriculture applications. Ann Agric Sci 65: 168–178.

