## Supplementary Figure S1 for "Induced systemic resistance by the root colonization of *Trichoderma atroviride* is independent from the chitin-mediated signaling pathway in Arabidopsis"

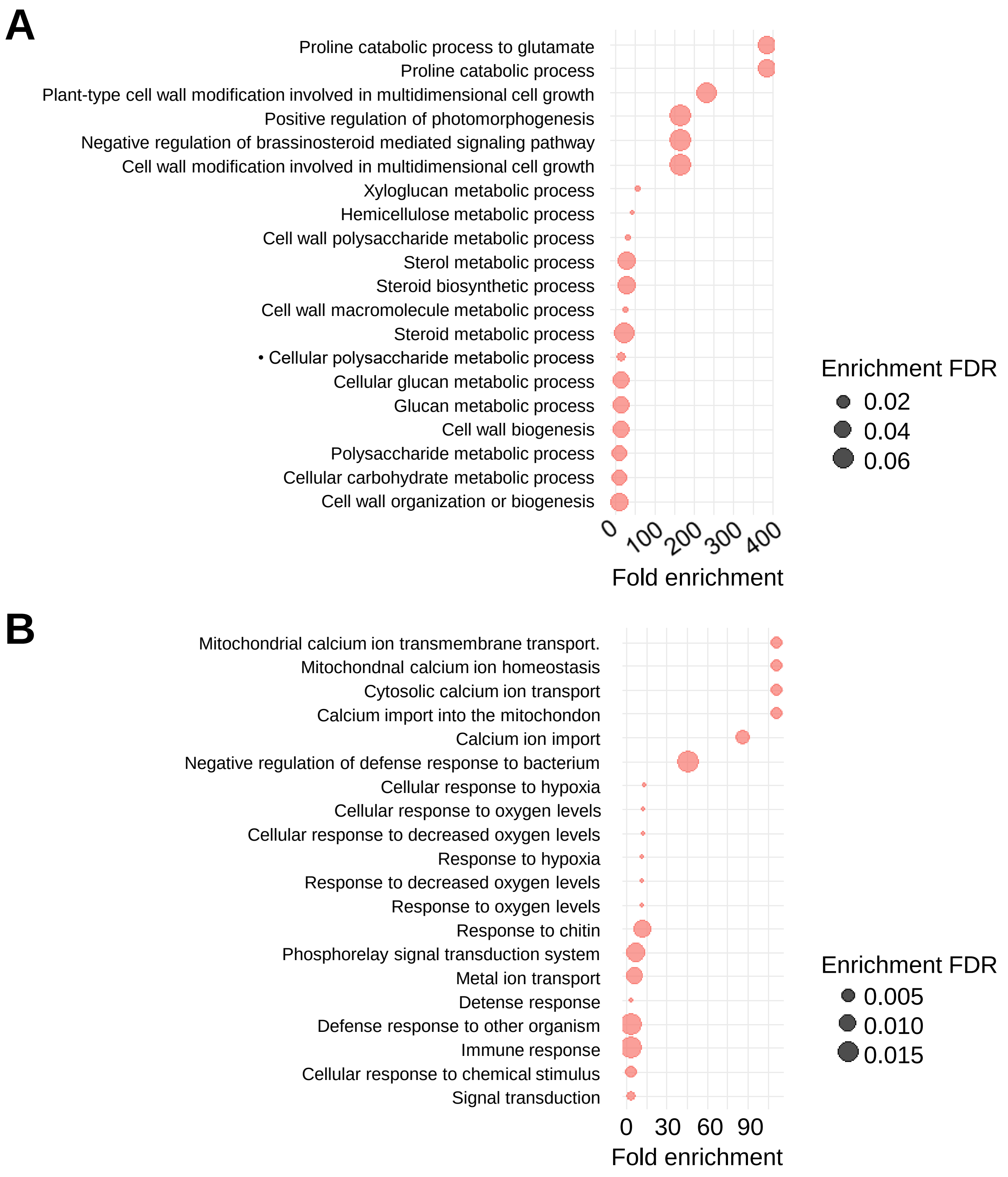


**Supplementary Fig. S1.**
GO enrichment analysis of the leaves of Arabidopsis seedlings treated with chitin. GO enrichment analysis of upregulated (A) and downregulated (B) DEGs regulated by chitin treated. The circle size indicates the FDR value. These figures showed the top 20 GO terms with the lowest FDR values in the biological process dataset.
